# A paired liver biopsy and plasma proteomics study reveals circulating biomarkers for alcohol-related liver disease

**DOI:** 10.1101/2020.10.16.337592

**Authors:** Lili Niu, Maja Thiele, Philipp E. Geyer, Ditlev Nytoft Rasmussen, Henry Emanuel Webel, Alberto Santos, Rajat Gupta, Florian Meier, Maximilian Strauss, Maria Kjaergaard, Katrine Lindvig, Suganya Jacobsen, Simon Rasmussen, Torben Hansen, Aleksander Krag, Matthias Mann

## Abstract

Existing tests for detecting liver fibrosis, inflammation and steatosis, three stages of liver disease that are still reversible are severely hampered by limited accuracy or invasive nature. Here, we present a paired liver-plasma proteomics approach to infer molecular pathophysiology and to identify biomarkers in a cross-sectional alcohol-related liver disease cohort of nearly 600 individuals. Metabolic functions were downregulated whereas fibrosis-associated signaling and novel immune responses were upregulated, but only half of tissue proteome changes were transmitted to the circulation. Machine learning models based on our biomarker panels outperformed existing tests, laying the foundation for a generic proteomic liver health assessment.

---

Liver disease has become one of the top causes of mortality worldwide, with non-alcoholic fatty liver disease (NAFLD) and alcohol-related liver disease (ALD) driving a rapid increase in incidence (Estes et al., 2018; Sheka et al., 2020; Tapper and Parikh, 2018). ALD is one of the most prevalent types of chronic liver disease worldwide, overtaking Hepatitis C virus infection in the United States (Cholankeril and Ahmed, 2018), and is the cause of more than half of all liver-related deaths; at least 500,000 fatalities every year (Pimpin et al., 2018; Sheron, 2016). ALD progresses through a range of histological lesions starting with alcohol-related fatty liver, to subclinical steatohepatitis (ASH) featuring hepatic inflammation, which drives fibrosis, and ultimately cirrhosis. These types of liver lesions can occur individually and concomitantly in the same patient to varying degrees, and often occur together with obesity, which further complicates the phenotypic spectrum of the disease (Harris et al., 2019). Nearly all individuals with chronic heavy alcohol consumption (>40 g of alcohol per day) develop fatty liver, 10 to 35% of these progress to ASH and 8-20% to cirrhosis (Seitz et al., 2018). However, the slow and generally asymptomatic nature of disease progression renders diagnosis at an early stage challenging. Moreover, accurate diagnosis of liver disease requires biopsy, a procedure that causes major complications in 1% of cases (Takyar et al., 2017). Ultrasound- and blood-based approaches are minimally invasive and focus on individual histological aspects of the disease, but has limited accuracy especially at early stages, thus severely reducing treatment options (Li et al., 2018). Therefore, there is a pressing need for minimally invasive diagnostic strategies for screening patients in at risk populations.

Biomarker discovery efforts have typically focused on individual biomolecules and have had a low acceptance rate in the clinic. Systems-wide studies, in contrast, have not connected changes in circulating levels and dysregulation in the diseased organ (Niu et al., 2019a). Given the high frequency of comorbid conditions in liver disease, such as chronic pancreatitis, cardiovascular disease, and cirrhotic cardiomyopathy (Jepsen, 2014; Keeffe, 1997), a connection in protein changes between liver and plasma would greatly help in evaluating their organ specificity and role in the disease (Veyel et al., 2020).

Recent advances in mass spectrometry (MS)-based proteomics have greatly extended its reach in biomedical and clinical research (Aebersold and Mann, 2016; Altelaar et al., 2013; Budayeva and Kirkpatrick, 2020). It is an increasingly powerful platform for specifically identifying and quantifying hundreds to thousands of proteins present in biological or clinical samples, making it highly suitable for identifying disease biomarkers. This is especially evident in the context of complex diseases, where focus on a single, or a handful of proteins is unlikely to provide accurate and reliable diagnosis. However, to be effective as a clinical biomarker discovery platform, MS-based proteomics has to be performed in a robust and accurate manner and applied to large patient cohorts. Recently our group has developed a plasma proteome profiling workflow and identified circulating proteins associated with NAFLD (Geyer et al., 2016; Niu et al., 2019b).

Here, we used MS-based proteomics to analyze paired liver tissue and plasma samples from a large cohort of patients and age- and gender-matched healthy volunteers with the goal of identifying circulating biomarkers for different pathological features of ALD. We demonstrated that liquid biopsies could be an effective replacement for more invasive liver ones. We observed that both liver and plasma proteomes undergo extensive remodeling during ALD, with fibrosis having the largest effect, followed by hepatic inflammation and steatosis. Our integrative analysis, cross-referencing liver and plasma proteomes, enabled us to use machine-learning to define plasma-based proteomics biomarker panels which outperformed existing liver tests in identifying early stages of liver fibrosis, inflammation and steatosis, as well as to predict future liver-related events. Finally, we will provide an interactive data visualization app built within the open-source Dash framework for data exploration (Supplementary Fig. 1).

## Results

### Framework for integrated tissue and plasma biomarker discovery in a large scale ALD cohort

We reasoned that pairwise correlations in protein levels between liver biopsy and plasma of the same individual would generate the data necessary to systematically correlate liver histology with pathophysiology-dependent changes in the plasma proteome (Fig. 1). We measured the liver and plasma proteomes using a data independent acquisition (DIA) strategy and a state of the art single-run workflow (Guo et al., 2015; Meier et al., 2018). We applied this framework to a cohort of 361 ALD patients (high risk group), making it the largest biopsy-verified ALD study (for baseline characteristics see (Thiele et al., 2016; Thiele et al., 2018)). Another cohort of 98 individuals with a history of excessive drinking but benign liver (according to FibroScan results) was also included (at risk group, no biopsy taken), as well as a healthy cohort of 137 individuals. In total, we performed MS-based proteomics on nearly 600 plasma samples and 79 liver biopsies spanning the full range of fibrosis grades (F0 to F4). Comprehensive clinical data was acquired from all participants, including the currently best performing non-invasive fibrosis tests: the ELF blood test (Enhanced Liver Fibrosis, (Guha et al., 2008)) for 380 of them, and 576 test results from transient elastography — an ultrasound-based technique for liver stiffness measurement. We also analyzed plasma proteome data using machine learning to construct plasma-derived ALD disease progression classifiers, and compared their performance to ELF, FibroScan and 11 other biomarkers.

**Fig. 1.**
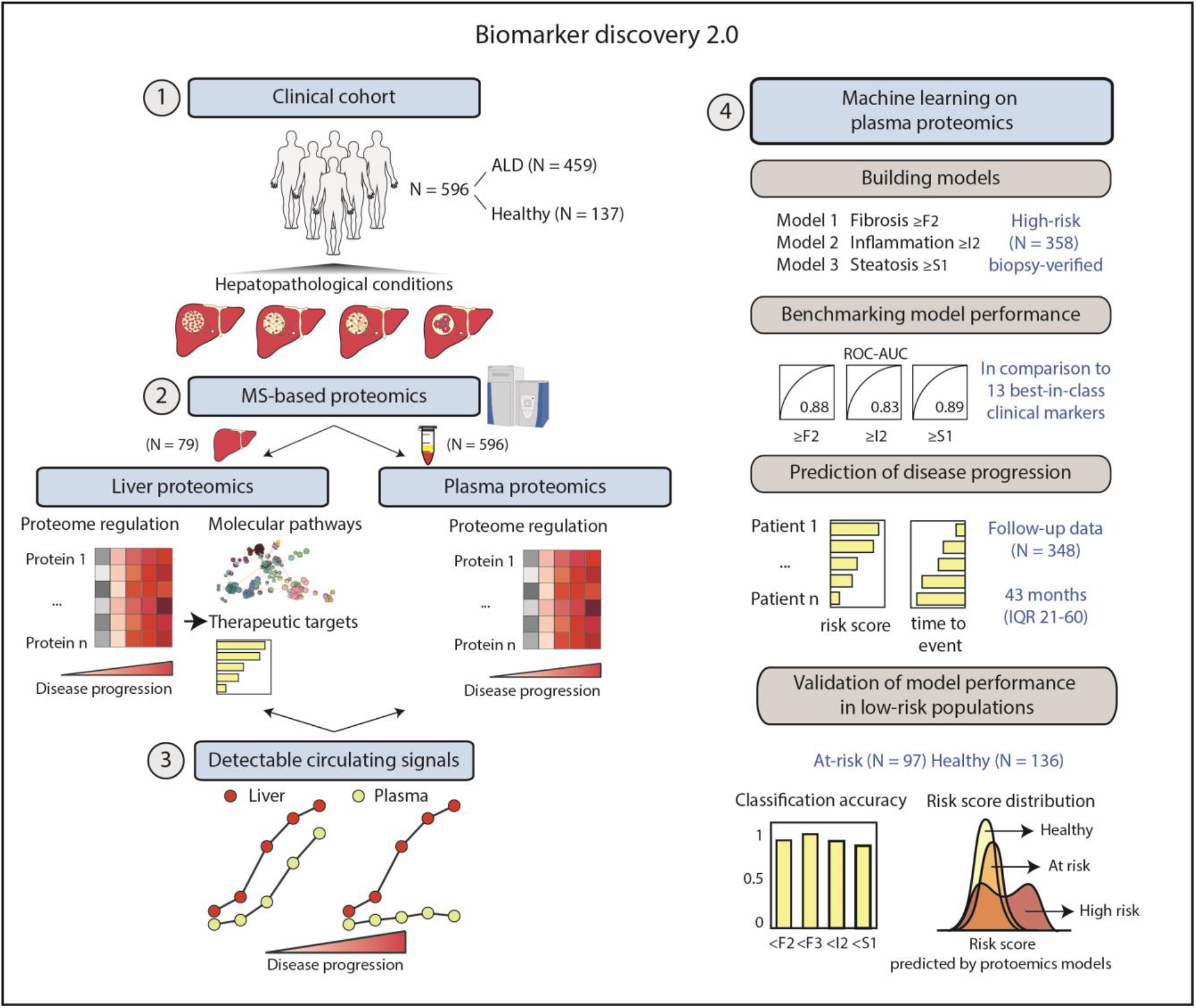
A framework for biomarker discovery in liver disease. In this framework, we applied high-throughput MS-based proteomics technology to profile paired liver and plasma samples from patients in a large clinically well-characterized cohort of ALD and matched healthy controls. Proteome regulation in the liver and plasma revealed changes at pathway and biological processes levels. Integrated liver and plasma proteomics helps to capture disease stage-relevant protein signatures in the bloodstream which are concordant with the liver. Lastly, we built a machine learning model to identify early stages of liver fibrosis, inflammation and steatosis.

### Impact of hepatic lesions on the liver tissue proteome

To maximize proteome depth and coverage from needle biopsies, we acquired liver proteomes of 79 patients in 100 min single-runs with an optimized DIA method (Supplementary Fig. 2). This was enabled by MaxQuant.Live (Wichmann et al., 2019) with a novel signal processing algorithm for fast LC-MS/MS cycle times (Grinfeld et al., 2017). Matching to a liver tissue library of more than 10,000 proteins yielded 5,515 total quantified proteins (Methods, Supplementary Fig. 3). Further filtering for proteins with less than 40% missing values across all samples resulted in a final set of 4,765 proteins with 95% data completeness. Differential protein expression analysis, controlled for common covariates, resulted in identification of 764 proteins significantly dysregulated in different stages of liver disease (fibrosis (701 proteins), inflammation (153 proteins) and steatosis (73 proteins)). Collectively, this accounts for 14% of the total quantified proteome (Fig. 2a, Supplementary Table 1). Fibrosis has by far the most dramatic effect on the liver proteome, followed by inflammation, with a large overlap between the two (86% of all proteins dysregulated in inflammation are also found to be dysregulated in fibrosis).

**Fig. 2.**
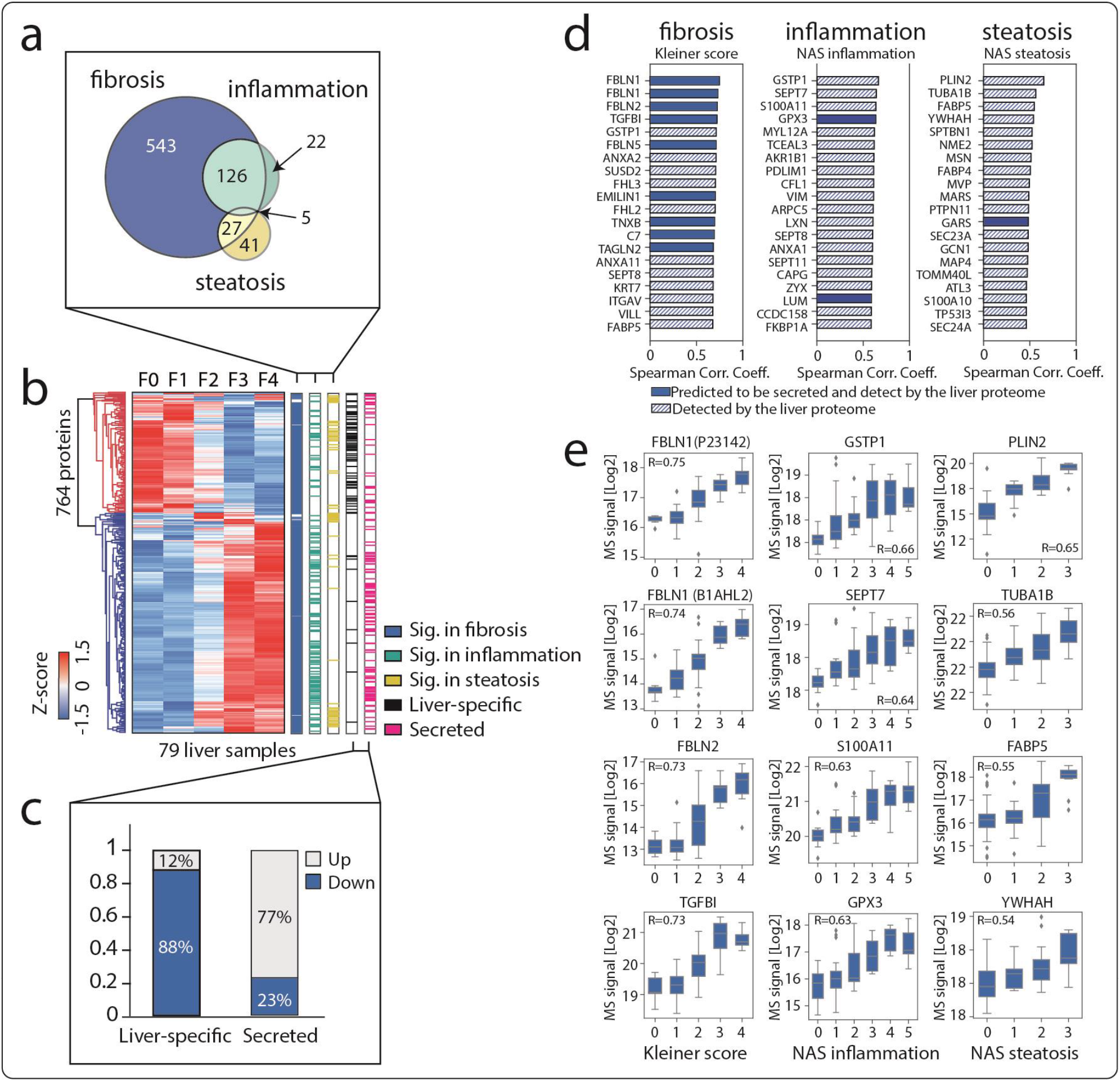
Liver proteome remodeling due to hepatic lesions. **a**. Venn diagram of the number of proteins in liver tissue significantly differentially abundant across stages of fibrosis, inflammation and steatosis, assessed by ANCOVA on 79 liver tissues followed by Benjamini-Hochberg correction for multiple hypothesis testing with a false discovery rate (FDR) of 0.05. **b**. Hierarchical clustering of all 764 dysregulated proteins that are significant in the ANCOVA. Row clustering was based on median log2-intensity after Z-score normalization across fibrosis stages F0-F4. Proteins significant across stages of fibrosis, inflammation and steatosis are color-coded as well as proteins categorized as ‘liver-specific’ and ‘secreted’ according to the Human Protein Atlas. Two major clusters of proteins that were down- and upregulated as degrees of fibrosis increased were color-coded in blue and red in the dendrogram, respectively. **c**. Fraction (%) of up- and downregulated ‘liver-specific’ and ‘secreted’ proteins. **d**. Top 20 proteins that correlate with the Kleiner score, NAS inflammation score, and NAS steatosis score, respectively. Proteins colored in blue are secreted according to the Protein Atlas. **e**. The distribution of log2-intensity values of top four correlating proteins to each histologic score. Number of replicates is in (Supplementary Fig. 4). The black line in the middle of the box is the median, the top and the bottom of the box represent the upper and lower quartile values of the data and the whiskers represent the upper and lower limits for considering outliers (Q3+1.5*IQR, Q1-1.5*IQR). IQR is the interquartile range (Q3–Q1).

The majority of dysregulated proteins across the fibrosis stages were found to be upregulated (65% (497 proteins) vs. 35% (267 proteins) downregulated; Figure 2b). We also grouped the dysregulated proteins based on the Human Protein Atlas annotation (Kampf et al., 2014), and found that 88% of the proteins annotated as ‘liver specific’ (Methods) were downregulated in liver tissue along increasing stages of fibrosis, while the effect on proteins annotated as ‘secreted’ was the opposite, with 77% of those being upregulated (Fig. 2c). This is similar to what we previously reported in plasma of non-alcoholic fatty liver disease (NAFLD), a related liver disease that exhibits pathophysiological similarities with ALD (Niu et al., 2019b).

Functional pathway annotation and enrichment analysis of the dysregulated liver proteome revealed that approximately 25% of the upregulated proteins belong to the immune system, specifically proteins involved in neutrophil degranulation (including the cell surface markers CD44 and CD63) and interleukin signaling pathways (Supplementary Table 2). Our results reflect the involvement of both innate and adaptive immunity in the pathogenesis of ALD at the molecular level. Signal transduction pathways were the second most highly over-represented category of proteins as fibrosis progresses (21% of total upregulated proteins), most notably receptor tyrosine kinase (RTK) signaling mediated by TGF-β, PDGF, MET, SCF-KIT and the insulin receptor, in agreement with existing literature that both TGF-β and PDGF play a role in fibrogenesis (Tsuchida and Friedman, 2017). Specifically, MAPK1, MAPK3, ROCK2 and AKT2, phosphatases PTPN6 and PTPN11, transcription factors STAT1, YAP2, and the small GTPases RAC1 and RAP1B were upregulated between fibrosis stage F0 to F4 (Supplementary Table 1). Signaling by Rho GTPase (40 proteins) and GPCR (23 proteins) were also significantly enriched in the upregulated proteins. Additionally, as expected based on their role in fibrosis, extracellular matrix (ECM) structural proteins and those involved in their biosynthesis, degradation, and post-translational modification were found to be significantly upregulated (46 proteins, including collagens type I, III, IV, V, VI, XII, XIV, fibronectin, laminin, lumican, perlecan, fibulins FBLN1, FBLN2, FBLN5, and the latent-transforming growth factor beta-binding proteins LTBP1, LTBP4; Supplementary Table 1). The proteins observed to be downregulated are largely comprised of metabolism-associated proteins (153 proteins, 57%), involved in metabolism of lipids, amino acids, xenobiotics, carbohydrates, steroids, cholesterol, vitamins and cofactors, as well as bile acids (Supplementary Table 3).

We also assessed correlation between measured proteomic changes and the widely used histological scores of liver pathology: the Kleiner fibrosis score (F0-4), the sum of NAS lobular inflammation and ballooning scores reflecting activity grade (I0-5), and the NAS steatosis score (S0-3) (Kleiner et al., 2005). We observed that 1,288 proteins significantly correlated with fibrosis stage, 951 proteins to the inflammatory activity, and 237 proteins to steatosis (Supplementary Table 4-6). Among the top 20 fibrosis-associated proteins ranked by correlation coefficient were prominent liver secreted proteins (Fig. 2d), with roles in cell-ECM interactions (TGFBI and EMILIN1, (Karsdal et al., 2015)), as well as proteins previously associated with hepatic fibrosis (Transgelin-2, (Molleken et al., 2009) and tenascin-X, (Kasprzycka et al., 2015)). Approximately 50% of the top 20 inflammation-correlated proteins are cytoskeletal proteins, indicating cellular structural changes in inflamed liver (Fig. 2d), while the top 20 hepatic steatosis-associated proteins include extracellular vesicle proteins reflecting changes in lipid transport. The four most significantly regulated proteins for each histological score are shown in Fig. 2e, with fibulins (FBLN1 and FBLN2) best correlated with the fibrosis score; glutathione metabolizing enzymes (GSTP1 and GPX3) and SEPT7, a protein with a known role in hepatocellular carcinoma (SEPT7) exhibited best correlation with the inflammation score. The lipid-droplet binding protein perilipin-2 and fatty acid-binding protein 4 (FABP4), an adipokine previously identified as a predictive marker for the progression from simple steatosis to NASH in patients with NAFLD (Coilly et al., 2019), exhibited some of the strongest correlation with the steatosis score (Fig. 2e). Taken together, proteomic analysis of the liver tissues representing different types of hepatic lesions shows that each has a unique proteomic signature, distinct from each other. Insights from functional annotation, pathway analysis and histological score correlation give confidence that the changes in protein abundance we measured in our proteomic experiments report on the status of ALD.

### Impact of hepatic lesions on the plasma proteome

Next we quantified and analyzed plasma proteomes from 360 biopsy-verified ALD patients, 137 healthy controls and 99 individuals who have a history of excess drinking but benign liver (Fig. 1; see Methods section for technical details, four samples were excluded from downstream analysis as they failed proteomics data quality criteria). Plasma proteomics is challenging because of the very high dynamic range, which we had previously addressed by multiple injections and the ‘BoxCar’ acquisition method (Meier et al., 2018; Niu et al., 2019b). To scale up to our cohort of 596 participants, we took advantage of the Evosep One, a novel LC system designed for clinical studies with faster analysis time, improved robustness and throughput (Bache et al., 2018). Furthermore, we applied a hybrid BoxCar (Meier et al., 2018) and DIA acquisition method (Ludwig et al., 2018), and a deconvolution method that doubles MS resolution (Grinfeld et al., 2017) (Methods).

Differential expression analysis revealed a total of 215 proteins significantly dysregulated in at least one comparison across histologic stages of fibrosis (88%), inflammation (62%) and steatosis (25%), underlying dramatic remodeling of the plasma proteome as a function of liver pathology (Fig. 3a, and Supplementary Table 7). Among these, 127 proteins were up- and 88 downregulated across fibrosis stages (Fig. 3b). The majority of dysregulated proteins overlapped between fibrosis and inflammation (Fig. 3a). Of the dysregulated proteins plasma proteins, including both proteins with a known role in liver fibrosis (type VI collagen (COL6A3), lumican, VWF and fibulin-1 (FBLN1)), and less established ones (ADAMTS like 4 protein (ADAMTSL4) and extracellular matrix protein 1 (ECM1)), as well as enzymes with roles in regulating extracellular environment (sulfhydryl oxidase 1 (QSOX1) and metalloproteases MMP2 and ANPEP) (Supplementary Table 9). Moreover, we confirmed that two proteins previously reported as upregulated in NAFLD (TGFBI and galectin 3-binding protein annotated as ‘liver specific’ or ‘secreted’, 81% and 56% (LGALS3BP)) (Niu et al., 2019b), were upregulated in were found to be downregulated, respectively (Fig. 3c). The downregulated cluster was mainly comprised of the complement and coagulation cascade and lipoprotein particles (Supplementary Table 8, Supplementary Fig. 5). Complement components C4A, C4B, C6, C8A, C8G as well as C4-binding protein, decreased in abundance by 15% to 42% between F0 and F4 (Supplementary Table 9), whereas C7 increased 3.5-fold, consistent with the literature (Niu et al., 2019b). Additionally, we observed downregulation of: (i) coagulation (co)factors and fibrinolytic proteins (prothrombin (F2), coagulation factor X, XI, XII, carboxypeptidase2 (CPB2), Protein C (PROC), Protein S (PROS1), antithrombin-III (SERPINC1) and alpha-2-antiplasmin (SERPINF2)); (ii) apolipoproteins (APOM, APOF, APOE, APOC4 and APOL1); and (iii) carrier proteins (albumin, retinol-binding protein, vitamin D-binding protein and hemopexin). Many of these changes are consistent with a dynamic interplay between fibrosis and steatosis that accompanies fibrosis progression. Interestingly, our mass spectrometric results of above mentioned clotting factors significantly correlated with values of the international normalized ratio (INR) opening up the possible substitution of the INR assay by plasma proteomics (Supplementary Fig. 6).

**Fig. 3.**
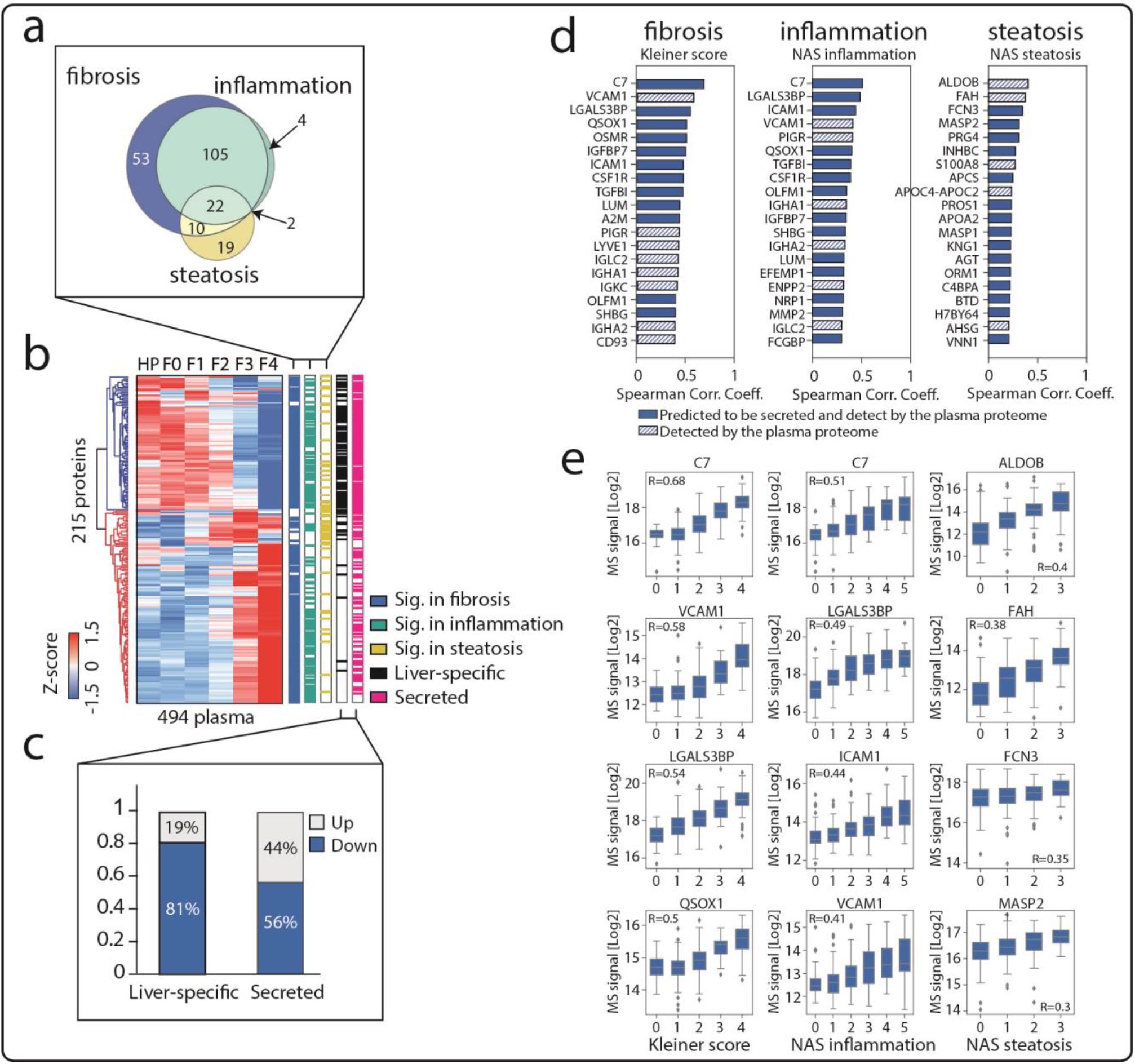
Remodeled plasma proteome due to hepatic lesions. **a**. Proteins in plasma significantly differentially abundant across stages of fibrosis, inflammation and steatosis as assessed by ANCOVA analysis on 494 plasma samples from the healthy controls (N=136) and the disease cohort with biopsy-verified histologic scores (N=358). ANCOVA significance level was corrected for multiple hypothesis testing by Benjamin-Hochberg at a false discovery rate of 0.05. **b**. Hierarchical clustering of the 215 significantly dysregulated plasma proteins from ANCOVA. Row clustering was based on median log2-intensity after Z-score normalization across fibrosis stages HP-F0-F4. Significant proteins across stages of fibrosis, inflammation and steatosis as well as ‘liver-specific’ and ‘secreted’ proteins according to the Human Protein Atlas were color-coded. Down- and upregulated as degrees of fibrosis increase were color-coded in blue and red in the dendrogram. **c**. Ratios of up- and down-regulating proteins of ‘liver-specific’ and ‘secreted’ proteins. **d**. Top 20 plasma proteins that correlate with the Kleiner score, NAS inflammation score, and NAS steatosis score. Proteins colored in blue are secreted proteins according to the Human Protein Atlas and the hatched boxes according to our plasma proteomics. **e**. Distribution of log2-intensity values of top four correlating proteins in plasma for each histologic score. Number of replicates is in (Supplementary Fig. 4).

Insulin-like growth factor-binding proteins (IGFBPs) IGFBP3, IGFALS, and IGFBP7, significantly and dramatically changed already at F3 stage (IGFBP3 and IGFALS decreased by 32% and 45% respectively, while IGFBP7 increased 2.1-fold). ECM-related proteins comprised a large group of upregulated ALD plasma. We also observed upregulation in: (i) cell adhesion molecules (ICAM1, VCAM1, integrin alpha-1 (ITGA1) and cadherin-5); and (ii) plasma immunoglobulins (IgA1-2, IgG1-4 and IgMs), as well as PIGR, a protein involved in transporting dimeric IgAs and polymeric IgMs (Mostov, 1994), which we previously identified as a promising marker for NAFLD (Niu et al., 2019b). PIGR was 2.6-fold upregulated in this cohort, together with J-chain (3-fold), a protein component responsible for PIGR binding to its ligand. Remarkably, PIGR was concordantly upregulated in the paired liver biopsies with an even larger increase (6.5-fold, Supplementary Table 1) further supporting its diagnostic value in liver disease.

Circulating levels of 79 proteins significantly correlated to fibrosis stage, 31 to inflammatory activity grade and five to the steatosis score (Fig. 3d; Supplementary Table 11-13). Complement component C7 (Spearman r = 0.68), VCAM1 (r = 0.58), LGALS3BP (r = 0.54) and QSOX1 (r = 0.5) had the highest correlation to fibrosis. Along with PIGR (5^th^ highest correlation coefficient), these are promising fibrosis marker candidates due to their roles in ECM remodeling and stage-dependent increase in the circulation. Fructose bisphosphate aldolase B (ALDOB), a key enzyme in aldolase metabolism, which we previously reported as a potential biomarker for NAFLD, had the highest correlation to steatosis (r = 0.4) (Fig. 3e). Collectively, this large-scale analysis demonstrates that liver status impacts the plasma proteome, resulting in specific, measurable changes in levels of subset of proteins with an interpretable role. Overall, this suggests that integrative analysis of liver and plasma proteomes of the ALD cohort may yield actionable circulating biomarker panels for each pathological lesion of ALD.

### Integration of liver and plasma proteomics

We analyzed liver and plasma proteomes further and identified overlapping proteins present in both datasets. Our analysis revealed a bi-modal pattern when matching protein levels in a 2-dimensional space (Fig. 4a-b). For one group of proteins, levels were largely correlated (‘diagonal cluster’), whereas for the other group relative abundance rank in liver was independent of that in the circulation (‘vertical cluster’, where each dot represents a protein, Fig. 4b). For example, liver enzymes ALDOB and carboxypeptidase D (CPD) exhibited about 1,000-fold difference in abundance in the liver, whereas their levels in plasma were very similar and low (Fig. 4c). In general, intracellular liver enzymes were part of the vertical cluster, including the well-known liver damage markers ALT (GPT), AST (GOT1), and gamma-glutamyl transferase (GGT, GGT1). Likewise, alcohol dehydrogenase 4 (ADH4), carboxylesterase 1 (CES1) and carbonyl reductase II (DCXR), all highly abundant in tissue (top 200, Supplementary Table 14) were detected in the plasma at very low levels - a million fold less than albumin, presumably reflecting tissue leakage.

**Fig. 4.**
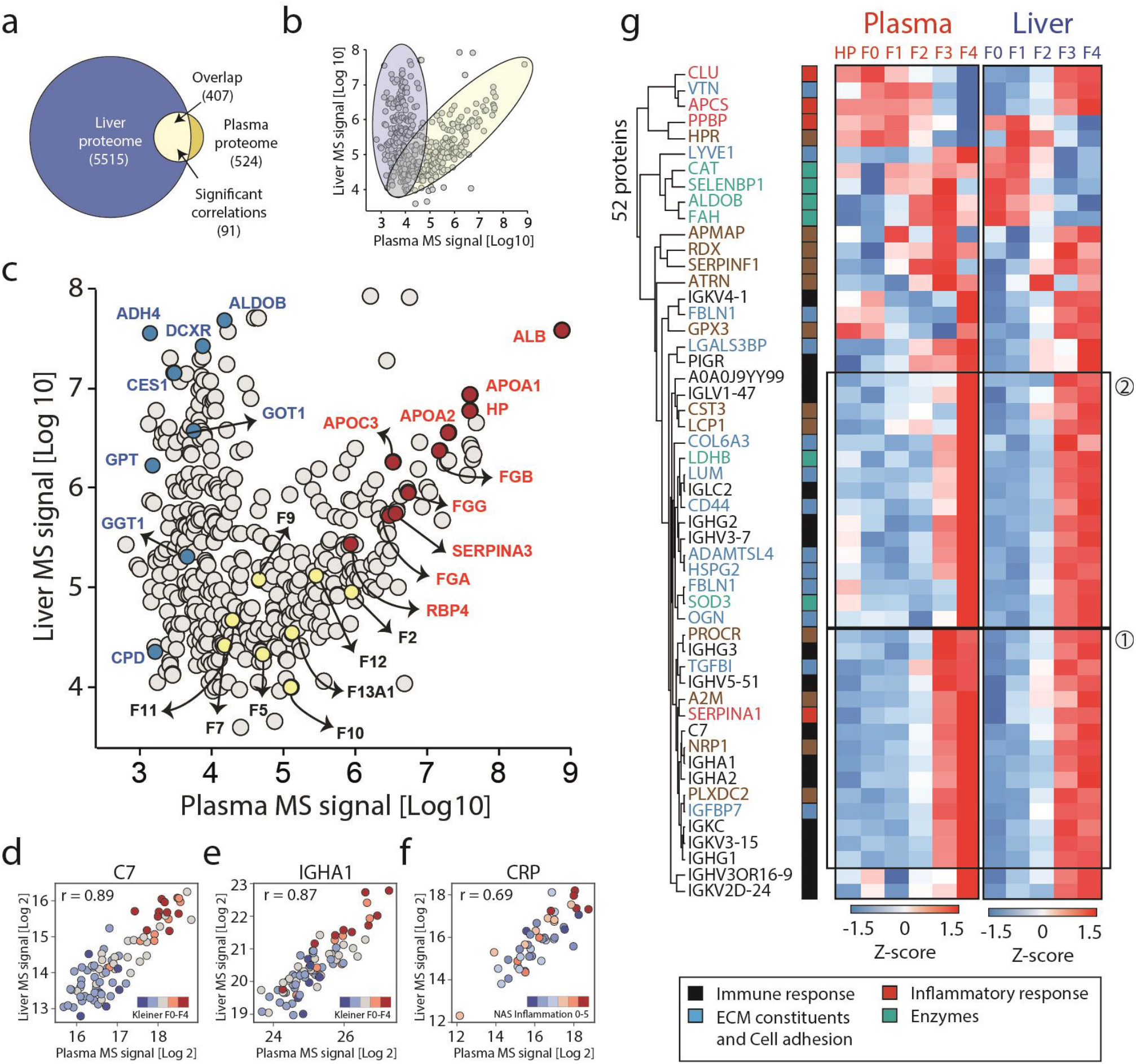
Liver-plasma proteome integration. **a**. Overlapping proteins between the liver- and plasma proteomes. Number of proteins that are significantly correlated between liver and plasma across patients is denoted (5% FDR and 0.3 minimum absolute Pearson correlation coefficient). **b**. Liver-plasma proteome abundance map showing median protein intensity (assessed by MS-intensity) in the liver as a function of that in the plasma. Proteins fall into a ‘diagonal cluster’ and a ‘vertical cluster’, characterized by dependence or independence of abundance rank in tissue vs. circulation, respectively. **c**. same as b) but color coding enzymes (blue), clotting factors (yellow) and known, functional plasma proteins related to the liver (darkred). **d-f**. Represented proteins significantly correlated in paired liver and plasma samples (C7, IGHA1 and CPR; Person correlation coefficient r indicated). Data points represent individual patients and are color-coded according to fibrosis score (Kleiner) or NAS inflammation score. Higher plasma and tissue abundances compared to other patients coincide with higher disease scores. **g**. Proteins co-regulated in the liver and plasma during disease progression. The dendogram shows the 52 significantly co-regulated proteins across histologic stages of liver fibrosis, inflammation and steatosis and their functional categorization as indicated in the bottom panel. The heat maps display their Z-scored median intensities across fibrosis stages within plasma (left) and liver (right). Clusters having ‘late’ (2) and ‘synced’ abundance patterns (1) in plasma compared to tissue are highlighted.

The diagonal cluster represents proteins that are secreted by the liver and act in the bloodstream, including abundant proteins like albumin, apolipoprotein A1, A2, C3, hepatoglobin (HP), SERPINA1 and fibrinogen subunits (FGA, FGB, FGG), as well as low abundant ones such as the coagulation factors F2, F9, F10, F11, F12 and F13A1 (Fig. 4c). Among commonly detected proteins, 91 had significant correlations between paired liver and plasma samples across the disease cohort (Pearson R up to 0.9 for C7 and IgA1, Supplementary Table 15). CRP levels in liver tissue and circulation also correlated highly (r = 0.69), which is of interest given the widespread use of this protein as a systemic risk marker for cardiovascular disease (Fig. 4d-f). PIGR’s value of 0.5 helps explain why both tissue and plasma levels correlated with disease severity.

Integrating the ANCOVA results of the liver and plasma proteomes resulted in 52 co-regulated proteins (Fig. 4g). These represent immune and inflammatory responses, extracellular matrix, cell adhesion molecules as well as intracellular enzymes and display distinct temporal patterns in the liver and plasma across fibrosis stages. The majority of them increased in abundance from fibrosis stage F0 to F4 in both liver and the plasma, with a few exceptions such as clusterin (CLU), vitronectin (VTN) and APCS. Apart from these monotonic patterns, ALDOB and fumarylacetoacetase (FAH) increased from F0 to F3 followed by a decrease in F4 in the plasma, but continuously decreased from F0 to F4 in the liver. Based on temporal profiles, we defined two clusters, both of which increased from F0 to F4 in tissue and circulation (Fig. 4g). Importantly, proteins of cluster 2 had a dramatic increase from F2 to F3 in the liver, but the signal was ‘delayed’ in the plasma – only dramatically increasing in F4 thus potentially representing late indicators of tissue damage. Conversely, proteins of cluster 1 had almost synchronized changes in the liver and the plasma. They included TGFBI, IGFBP7 and C7, all promising fibrosis markers that have been previously reported by us and others (Martínez-Castillo et al., 2020; Niu et al., 2019b).

### Machine learning models to detect early-stage fibrosis, inflammation and steatosis

To investigate whether plasma proteomics can identify early stage fibrosis (200 controls in F0-1, 160 cases in F2-4), we built a logistic regression model using a selected panel of a total of 14 proteins (Methods, Supplementary Table 16). Among these proteins, nine (64%) were co-regulated in plasma and liver, including IGHA2, HPR, PIGR, APCS, C7, LGALS3BP, TGFBI, IGFBP7 and LYVE1, of which four belong to the ‘synced’ signal pattern including IGHA1, IGFBP7, C7 and TGFBI (Fig. 4g). We compared the proteomic model against eight best-in-class clinical tests: transient elastography (TE), 2D-shear wave elastography (SWE), the ELF blood test, the FIB-4 Index, the FibroTest (FT), the APRI (AST to platelet ratio index), the P3NP and the Forns index (Thiele et al., 2016; Thiele et al., 2018). We used both clinically established cut-offs and logistic regression determined cut-offs to perform cross-validation (see Methods for details), and present the mean model performance to account for various missing values across clinical tests and the lack of an external validation dataset (Fig. 5). The 14-protein panel for predicting significant fibrosis (≥F2) resulted in a mean area under the receiver operator curve (AUROC) of 0.88 (95% CI: 0.82-0.94), F1 score of 0.82 (95% CI: 0.74-0.90), and a balanced accuracy of 0.81 (95% CI: 0.72-0.90) (Fig. 5a-d, Supplementary Table 17). Overall the proteomics model obtains the best combination of precision and recall as can be seen by the F1-score and balanced accuracy. The clinically defined ELF and FT markers obtain substantially higher precision or recall, but fail to balance both, leading to an inferior performance in the two other metrics. A final prediction model on a new random split demonstrated that the proteomic model was significantly superior to logistic regression models using either the APRI or the FIB-4 Index (p < 0.05), and equally as good as the others as shown by pair-wise DeLong’s tests between models (Supplementary Table 18). We similarly derived a nine-protein panel (Supplementary Table 16) for predicting early stage hepatic inflammation (sum of NAS lobular inflammation and NAS ballooning ≥ 2) (153 controls, 189 cases), which had an AUROC of 0.83 (95% CI: 0.74-0.92), F1 score of 0.77 (95% CI: 0.66-0.89) and balanced accuracy of 0.76 (95% CI: 0.65-0.87) (Fig. e-h, Supplementary Table 19). The F1 and balanced accuracy mean scores were higher than any of the other clinical comparators including the AST/ALT ratio (AAR), and the serum cytokeratin 18 epitopes M30 and M65 (Methods). The AAR test had the highest precision, but failed to capture many cases. DeLong’s test on the final model indicated that the proteomicFthi model was significantly superior to all other comparators (p < 0.05) (Supplementary Table 20). It selected LGALS3BP, C7, PIGR, IGFBP7, QSOX1, CLU, FBLN1, ICAM1 and SEPIND1 as features for the inflammation panel, six of which were co-regulated in plasma and liver (Fig. 4g).

**Fig. 5.**
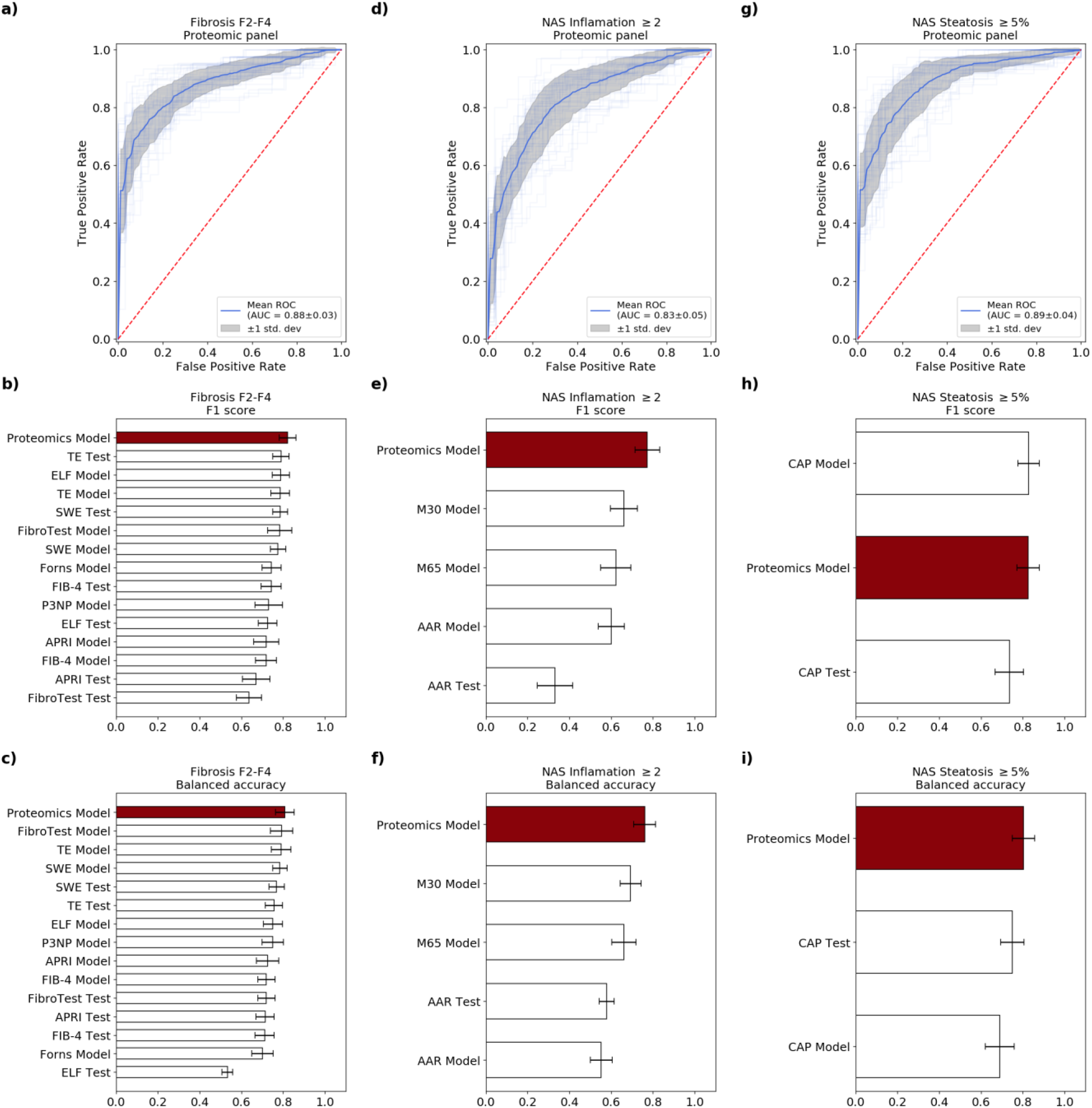
Prediction models based on the plasma proteome for biopsy-verified fibrosis, inflammation and steatosis. **a, d, g**. Receiver-operating characteristic (ROC) curve and corresponding area under the curve (AUC) statistics in 5-fold cross-validation repeated for 10 times of a protein panel-based logistic regression model for detecting significant liver fibrosis (F2-4) (a), hepatic inflammation (NAS I2-I5) and steatosis (NAS S1-S3) (g). **b-c, e-f, h-i**. Averaged AUROC, F1 score and balanced accuracy in cross-validation of the above-mentioned logistic regression models in comparison to best-in-class existing markers for fibrosis (b-c), inflammation (e-f) and steatosis (h-i). Performance of existing markers were calculated based on either their established clinical cut-offs and machine learning cut-offs.

Finally, a 28-protein panel-based model for predicting presence of any steatosis (NAS steatosis score ≥ 1) performed equally well as the controlled attenuation parameter (CAP), an ultrasonic measurement based on FibroScan for the detection of hepatic steatosis (Supplementary Table 21-22). Remarkably, the feature selection algorithm also picked ALDOB, a protein we previously identified as candidate marker for NAFLD, as well as lipid transport proteins APOC3 and apolipoprotein(a) (LPA), the latter being the main constituent of lipoprotein(a), a known risk factor for developing vascular disease. Taken together, these data strongly support the potential clinical value of plasma proteomics derived panels of proteins for detection of fibrosis, inflammation and steatosis simultaneously and for connecting them to their cellular and for tissue origins in ALD, and potentially NAFLD.

### ALD plasma biomarker model accuracy for excluding liver injury in healthy and at risk populations

Predicting risk scores for individuals in the healthy and at risk cohort with the final proteomic models, excluded significant fibrosis, advanced fibrosis and mild inflammation in the healthy, matched control population with accuracies of 95%, 100% and 96%, respectively (Supplementary Table 23). Thus, proteomics combined with machine learning appears to be able to rule out these conditions in healthy populations. Around 18% of the healthy individuals were classified as steatosis positive. However, since they were matched for BMI with the ALD population, and simple steatosis was not an exclusion criterion, clinical missassignment is possible. Supporting this, 60% of those who were classed as steatosis positive from the proteomics score had a CAP value above 290, the cut-off for fatty liver. This implies an even higher accuracy of the proteomics score for excluding steatosis. In the at risk population of patients with excess use of alcohol, but a benign liver assessed by FibroScan, 90% and 86% of individuals were predicted not to have significant fibrosis and inflammation, respectively. Individuals in this cohort did not meet the criteria of taking a liver biopsy based on the FibroScan results. This likely led to some under-diagnosis at the time of inclusion, explaining some of the remaining false-positives in our model.

To investigate if our proteomic marker panels predict future liver-related events, we followed 348 patients with a valid liver biopsy for a median of 43 months (IQR 21-60) after their study inclusion, using electronic patient files and Danish national registries to evaluate time to first liver related event. Seventy of them experienced a liver-related event during the follow-up period. To investigate the prognostic power of our proteomic marker panels, we computed their Harrell’s C-index and compared them with existing non-invasive markers and the liver histological lesions as reference (Methods). For significant fibrosis and moderate hepatic inflammation the C-index was 0.895 and 0.885, demonstrating that our proteomics marker panels for fibrosis and inflammation are highly accurate measures for predicting disease progression (Fig. 6c). Among existing tests, the score for TE was nearly as good, whereas the best serum markers (ELF) had a Harrell’s C inferior to three of our proteomic marker panels. Collectively, we demonstrate that the plasma biomarker panels that we constructed based on deep quantitative proteomic analysis offer a reliable, minimally invasive strategy for diagnosing, staging and predicting progression ALD.

**Fig. 6.**
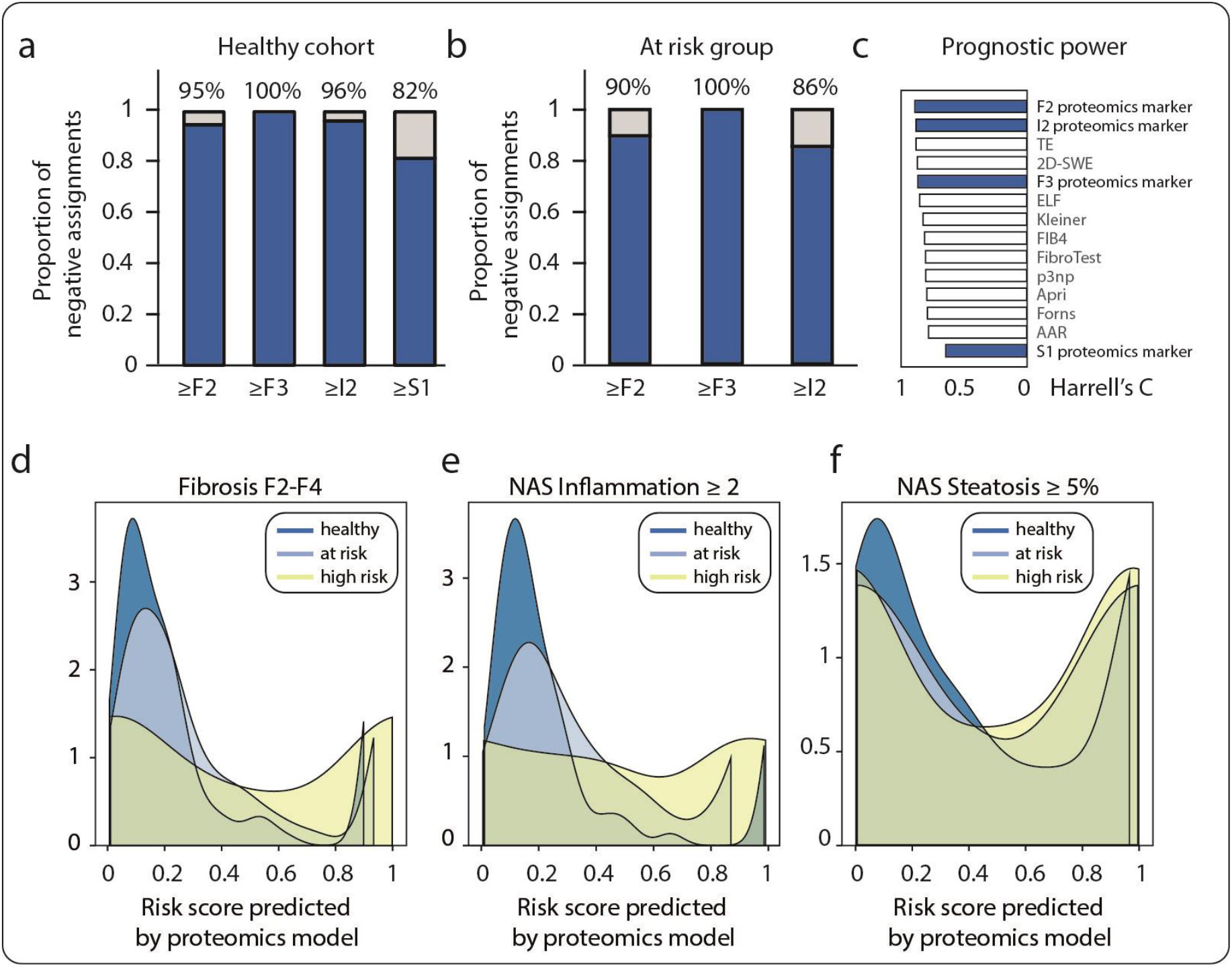
Additional validation of model performance and distribution of risk scores predicted by proteomics models. **a**. Percentage accuracy of the proteomics models for excluding significant fibrosis, advanced fibrosis, moderate inflammation and steatosis in the healthy cohort. **b**. Accuracy of the proteomics models for excluding significant fibrosis, advanced fibrosis and moderate inflammation in the at risk population. **c**. Harrell’s C-index of proteomics models and clinical tests in predicting patient outcome, ranked from the highest to lowest. **d-f**. Kernel density estimate plots showing the distribution of risk scores for significant fibrosis (c), moderate inflammation (d), and steatosis (d) predicted by respective proteomics models in the healthy, at risk- and high risk populations.

## Discussion

This work represents the largest and most detailed characterization of proteome changes in liver disease. Our analysis revealed that ALD progression is accompanied by dramatic proteome remodeling in both liver and the plasma, with fibrosis affecting the largest number of dysregulated proteins, followed by hepatic inflammation and steatosis. By integrating liver and plasma proteomics, together with clinical data on patients and healthy controls in our cohort we were able to define plasma biomarker panels for different hepatic lesions of ALD. When combined with machine learning, the plasma biomarker panels yielded predictive models that either performed as well as existing diagnostic strategies, or outperformed them, including in the context of predicting risk of future adverse liver events. Our study indicates that screening high risk individuals by proteomics could predict disease progression from an asymptomatic stage to symptomatic, therefore informing treatment options and intervention opportunities. Moreover, given the minimally invasive nature of plasma sample collection, proteomic screens could be performed with frequency not currently feasible, hence significantly aide the clinical decision-making process. Overall, our results suggest feasibility of developing liquid biopsies for a combined diagnosis of fibrosis, inflammation and steatosis, in addition to monitoring and prognostication. Confirmatory trials are needed to independently validate the approach.

Importantly, ALD and NAFLD share common histological presentations, therefore discovery of commonly regulated proteins could help develop a generic blood test for the detection of hepatic lesions irrespective of the initial cause. Notably, comparing our results to our previous, smaller NAFLD cohort (Niu et al., 2019b), we found PIGR, ALDOB, and LGALS3BP to be commonly upregulated with corresponding, disease-stage dependent changes in the paired liver tissue. Remarkably, PIGR, which emerged as a potential biomarker in the NAFLD study, was among the top 20 highest upregulated proteins in plasma between fibrosis stages F0 and F4 (2.6-fold), and the fifth highest in liver (6.5-fold). Taking the two studies together, PIGR emerges as a promising indicator for the presence and severity of fibrosis and inflammation, whereas LGALS3BP and ALDOB appear to indicate fibrosis and steatosis, respectively. Concordantly, these three proteins were selected by our machine learning algorithms in an unbiased way for predicting these lesions. We expect that proteomics study of larger NAFLD cohorts, enriched with additional clinical data conducted in the manner similar to the one described here will likely result in discovery of additional common biomarkers for NAFLD and ALD. More generally, a comprehensive study encompassing the major liver diseases could provide a set of common and distinguishing biomarkers in the future.

In addition to diagnostic power, our results also have major implications for the development of potential ALD and NAFLD treatment options. For example, an astonishing 20% of upregulated tissue proteins were associated with signal transduction pathways such as the receptor tyrosine kinase (RTK), including many kinases with available inhibitors in different stages of pre-clinical and clinical development. This may suggest a potential for targeting RTK signaling in the context of ALD. TGF-β signaling pathway represents a similar opportunity, given its function as the most potent driver of fibrogenesis and extensive existing drug development efforts (Beyer and Distler, 2013; Fabregat et al., 2016). Furthermore, proteomics implicated some less studied signaling pathways in ALD, including G-protein coupled receptors (GPCRs), widely considered to be druggable. Thus our results suggest additional targeting opportunities for developing pharmacological agents for ALD treatment.

Overall, here we outline a generally applicable framework for discovery and validation of plasma-based biomarker panels for diagnosing complex diseases, such as ALD. We highlight that availability of clinical samples and health record data from a large patient cohort, including paired tissue histology and plasma samples, as well access to paired samples from healthy individuals are critical for discovery of robust, predictive biomarker panels. We expect that combining quantitative MS-based proteomics of tissue of origin and paired patient plasma in other systems may lead to circulating biomarkers’ discovery in other diseases, where accurate diagnosis is currently based on biopsies requiring invasive procedures, resulting in a more widespread availability and application of less invasive liquid biopsies. With ongoing improvements in technology, we expect an increase in model performance. We envision that targeted or ‘global targeted’ (Wichmann et al., 2019) MS-based assays could be developed that retain the full specificity of this technology. An additional benefit of plasma proteomic profiling is its generic nature, meaning that it provides additional information apart from the targeted panel. For instance, bleeding and clotting abnormalities could be assessed in patients with liver disease as demonstrated by the correlation of the clotting factors to INR values in this study. A further advantage is that only a single technology needs to be applied.

## Materials and methods

### Ethical approval

The study protocol was approved by ethics committee for the Region of Southern Denmark (ethical IDs: S-20160006G, S-20120071, S-20160021, S-20170087) and registered with the Danish Data Protection Agency (13/8204, 16/3492), also with Odense Patient Data Exploratory Network under study identification number OP_040 (open.rsyd.dk/OpenProjects/da/openProjectList.jsp). The study was conducted according to the principles of the Declaration of Helsinki, and oral and written informed consent was obtained from all participants.

### Participant recruitment and clinical data collection

For the healthy and ALD cohorts, participant recruitment protocol including inclusion and exclusion criteria, setting and locations where data were collected can be found in the original report (Thiele et al., 2016; Thiele et al., 2018; Trošt et al., 2020). The same applies for sample- and clinical data collection procedure, detailed description of the cohort characteristics and anything related to the original clinical study design. The study population in the ALD cohort has been previously published for the evaluation of the studied fibrosis tests. Cut-offs for these clinical tests when compared with proteomics models can be found in Supplementary Table 27.

### Plasma proteome sample preparation

Plasma samples were prepared with a modified protocol based on previously published Plasma Profiling pipeline on an automated liquid handling system (Agilent Bravo) in a 96-well plate format(Geyer et al., 2016). In brief, proteins were denatured, reduced, alkylated and enzymatically cleaved into peptides. Resulting digestion mixture were loaded on Evotips (Bache et al., 2018).

In detail:

1. Transfer 5 μl of blood plasma sample into an Eppendorf 96-well plate on an Agilent Bravo liquid handling system (Plasma plate).
2. Make a 1:10 dilution by adding 45 μl of lysis buffer (10mM TCEP, 40mM CAA, 100mM Tris, pH8.5) into each well of the Plasma plate, mix thoroughly by pipetting 50 times up and down for a volume of 40 μl, and centrifuge the plate up to 300 x g.
3. Pipet 20 μl of 10-fold diluted plasma into a new 96-well plate (Digestion plate).
4. Heat the plate at 95°C for 10 min.
5. Move the Digestion plate to room temperature and cool it down for 5 min. Meanwhile prepare fresh trypsin/LysC mix in 0.05 (μg/μl) (total volume calculated by 20 μl per sample, 1:100 micrograms of enzyme to micrograms of protein).
6. Add 20 μl of trypsin/LysC mix into each well to a final volume of 40 μl.
7. Heat the Digestion plate at 37°C for 2h (enzymatic digestion).
8. Quench the reaction by adding 64 μl of 0.2% TFA, mix thoroughly by pipetting 20 times up and down. The digestion mixture can be frozen and stored at this stage.
9. Load 1μg digestion mixture onto a disposable Evotip C18 trap column (Evosep Biosystems, Denmark) according to the manufacturer’s instructions. Briefly, Evotips were wetted with 2-propanol, equilibrated with 0.1% formic acid, and then loaded using centrifugal force at 1500 g with a 3D-printed centrifugal block. Evotips were washed with 0.1% formic acid and then added 200 μl of 0.1% formic acid to prevent drying.

### Liver proteome sample preparation

Snap-frozen liver biopsies were cryo-pulverized in a Covaris cryoPREP Dry Pulverizer and collected in a glass tube. Approximately 1mg (in some cases less than 1mg) of tissue powder was transferred to an Eppendorf tube, adding 150 μl of SDC reduction and alkylation buffer (PreOmics GmbH). The homogenate were then heated at 95°C for 10 min, vortexed at 1200 rpm on a thermos mixer (Eppendorf) to denature proteins, subsequently sonicated using a water bath sonicator (Diagenode Bioruptor^®^, Liège, Belgium) at full power for 30 cycles with 30s intervals and a second round sonication using the Covaris Adaptive Focused Acoustics (AFA) sonication system (Covaris, USA).

Protein content was determined by Tryptophan assay and a volume containing 50 μg of protein was digested overnight with trypsin and LysC (1:50, μg of enzyme: μg of protein) at 37°C, 1200rpm on a thermos mixer. Digestion mixture was acidified to a final concentration of 0.1% trifluoroacetic acid (TFA) to quench the digestion reaction. Peptide concentration was estimated using Nanodrop and 20 μg of peptide mixture was purified by solid phase extraction in a Stage-Tip format (SDB-RPS material, two 14-gauge plugs), washed with isopropanol/1% TFA and 0.2% TFA (200 μl each, centrifuge at 1500 x g with 3D-printed centrifuge block). Peptides were eluted with 60 μl of 80% acetonitrile/1% ammonia and dried at 60°C using a SpeedVac centrifuge (Eppendorf, Concentrator plus). Dried peptides were dissolved and sonicated in 5% acetonitrile/0.1% TFA. Peptide concentration was measured using Nanodrop, and 500 ng of purified peptides were injected for LC-MS-MS analysis.

### Liquid chromatography and mass spectrometry (LC-MS)

The acquisition of samples was randomized to avoid bias. In single-shot plasma proteome analysis, peptide mixture was partially eluted from Evotips with less than 35% acetonitrile and analyzed with an Evosep One liquid chromatography (LC) system (Evosep Biosystems, Denmark (Bache et al., 2018)) coupled online to a hybrid quadrupole Oribitrap mass spectrometer (Q Exactive HF-X). Eluted peptides were separated at 60 °C on a 15 cm long column (150 μm inner diameter packed with 3 μm Reprosil-Pur C18 beads (Dr. Maisch, Ammerbuch, Germany)) in a standard preset gradient method (44min, 30 samples per day), and electrosprayed from a laser-pulled silica emitter tip at 2.4 kV. Data was acquired in BoxCar/DIA mode (BoxCar/DIA window setting see Supplenmentary Table 26), enabled by MaxQuant.Live (Wichmann et al., 2019) in which the scan protocol was defined. Each acquisition cycle was consisted of a survey scan (automatic gain control target (AGC) of 3e6 or 100ms injection time), two BoxCar scans (12 windows each at 300 - 1,650 m/z, AGC of 1e6/120ms) both at a resolution of 60, 000 at m/z 200, followed by 29 DIA cycles (AGC of 3e6/54ms at range 300-1650 m/z). HCD fragmentation was set to normalized collision energy of 27%. In all scans, PhiSDM (Grinfeld et al., 2017) was enabled with 100 iterations, spectra type was set to centroid.

In single-shot liver proteome analysis, purified peptides were measured using LC-MS instrumentation consisting of an EASY-nLC 1200 system (Thermo Fisher Scientific, San Jose, CA) interfaced on-line with a Q Exactive HF-X Orbitrap (Thermo Fisher Scientific, Bremen, Germany). Peptides were separated on 42.5 cm HPLC-columns (ID: 75 μm; in-house packed into the tip with ReproSil-Pur C18-AQ 1.9 μm resin (Dr. Maisch GmbH)). For each LC-MS/MS analysis, around 0.5 μg peptides were injected for the 100 min gradients. Peptides were loaded in buffer A (0.1% formic acid) and eluted with a linear 82 min gradient of 3-23% of buffer B (0.1% formic acid, 80% (v/v) acetonitrile), followed by a 8 min increase to 40% of buffer B. The gradients then increased to 98% of buffer B within 6 min, which was kept for 4 min. Flow rates were kept at 350 nl/min. Re-equilibration was done for 4 μl of 0.1% buffer A at a pressure of 980 bar. Column temperature was kept at 60 °C using an integrated column oven (PRSO-V2, Sonation, Biberach, Germany). Data were acquired using an optimized DIA method, enabled by MaxQuant.Live (Wichmann et al., 2019) in which the scan protocol was defined. Each acquisition cycle was consisted of a survey scan at resolution of 60,000 with an automatic gain control target (AGC) of 3e6/100ms injection time (IT), followed by 66 DIA cycles (Supplementary Table 26) at resolution of 15,000 with an AGC of 3e6/22ms IT at range 300-1650 m/z). HCD fragmentation was set to normalized collision energy of 27%. In all scans, PhiSDM (Grinfeld et al., 2017) was enabled with 100 iterations, spectra type was set to centroid.

### Mass spectrometric data analysis

BoxCar/DIA hybrid spectra in the plasma dataset and DIA spectra in the liver biopsy dataset were analyzed with Spectronaut v13. The default settings were used unless else noted. Data filtering was set to ‘Qvalue’. ‘Cross run normalization’ was enabled with strategy of ‘local normalization’ based on rows with ‘Qvalue complete’. The false discovery rate (FDR) was set to 1% at peptide precursor level and 1% at protein level. A previously generated deep fractionated plasma DDA library (Geyer et al., 2016; Niu et al., 2019b) and liver DDA library were used in the targeted analysis of DIA data for plasma and liver datasets against the human UniProt fasta database (January 2018, 21,007 canonical and 72,792 additional sequences).

### Pre-processing of liver and plasma proteomics datasets

Proteome datasets of the liver and plasma were filtered for 60% valid values across all samples (proteins with >40% missing values were excluded from downstream statistical analysis), with the remaining missing values imputed by drawing random samples from a normal distribution with downshifted mean by 1.8 standard deviation (SD) and scaled SD (0.3) relative to that of abundance distribution of the corresponding protein across all samples. Specifically in total, 524 proteins were quantified in the plasma proteomes, filtering for 60% valid values across all samples resulted in a dataset of 304 proteins with a data completeness of 95%. Assessed on seven quality control samples median workflow coefficient of variation (CV) was 23% across a one-month period (Supplementary Fig. 3). Liver proteome was pre-processed in the same way and more details were provided under the Results section (Supplementary Fig. 3).

### Differential expression analysis

Differentially expressed proteins were determined by analysis of covariance (ANCOVA) controlling for the common covariates age, BMI, sex, and abstinence status at inclusion. We also controlled for the effects of steatosis when assessing the effect of fibrosis and inflammation on the liver proteome, and vice versa. Histological stages include a five-grade fibrosis score (F0-F4, denoted as Kleiner), a six-grade inflammation score (NAFLD Activity Score, NAS_I0-I5) combining lobular inflammation and ballooning, and a four-grade steatosis score (NAS_S0-S3). A Python script based on an open source statistical package pingouin.ancova was developed to handle ANCOVA in proteomics data and control for multiple hypothesis testing. A protein was considered significantly differentially expressed across a given condition if the ANCOVA-derived FDR-adjusted P-value by Benjamini-Hochberg was below 0.05. Significant proteins and corresponding P-values are provided in Supplementary Table 1 and 7. Differential expression across disease stage is presented as a heatmap generated by the Perseus computational software (version 1.6.5.0) (Tyanova et al., 2016).

### Proteome correlation to histology scores

Spearman partial correlation controlling for the same covariates as ANCOVA was performed to assess protein-histology score correlation. A correlation was considered significant if the FDR-adjusted P-value by Benjamini-Hochberg was below 0.05 and the absolute value of correlation coefficient r equals to or is larger than 0.3. A python script based on an open source statistical package pingouin.partial_corr was developed to handle partial correlation in proteomics data and control for multiple hypothesis testing. Significant proteins and corresponding P-values, Spearman correlation coefficients were provided in Supplementary Table 4-6 and 11-13. Protein expression in dependence of disease stage was presented as bar plots generated in Jupyter notebook environment.

### Pairwise liver-plasma proteome correlation

Pair-wise correlation was performed to assess the correlation between paired liver biopsy and plasma across the patient cohort. Significance level was controlled at an FDR-adjuster P-value by Benjamini-Hochberg of below 0.05 and an absolute value of correlation coefficient r larger than 0.3. A python script based on an open source statistical package pingouin.pairwise_corr was developed to handle pairwise correlation in proteomics data and control for multiple hypothesis testing. Significant proteins and corresponding P-values, Pearson correlation coefficients and annotations for human protein atlas were provided in Supplementary Table 15. Selected significantly correlating proteins were presented as scatter plots with MS signal in liver biopsy as a function of that in plasma.

### Functional Annotation and Enrichment Analysis

Ontology Enrichment Analysis in the liver proteomics dataset was performed with ClueGo, a plug-in app in Cytoscape, with default settings. Customed reference set which contains 4,651 unique genes (quantified in this study) was used in Fisher’s exact test. Term significance was corrected by Benjamini-Hochberg with a FDR of below 1%. Both GO term fusion and grouping were activated. The equivalent in the plasma proteomics dataset was performed in the Perseus software. Significantly enriched Reactome and GO terms and associated proteins were provided in Supplementary Table 2-3 and 8-10.

Liver-specific proteins were annotated according to the Human Protein Atlas (HPA), which defines ‘liver enriched’, ‘group enriched’, and ‘liver enhanced’ proteins with at least five-times higher mRNA levels in liver compared to all other tissues, at least five-times higher mRNA levels in a group of two–seven tissues compared to the rest, and at least five-times higher mRNA levels in the liver compared to average levels in all tissues, respectively.

### Machine learning models

The machine learning part of this manuscript is conducted with the intention to identify biomarker panels for identifying different types of hepatic lesions. A graphic representation of the overall machine learning workflow including classification targets, strategies for feature selection and model performance evaluation, assessment for prognostic power and validation in low-incidence populations can be found in supplementary Fig. 7.

More specifically, 360 patients had liver biopsy-verified stages of fibrosis (360), inflammation (352) and steatosis (352) as well as clinical data to varying degrees’ of missing values from best-in-class clinical tests, e.g. 331 SWE, 264 M65 or 199 CAP. Two of these patients were excluded due to insufficient proteome depth quantified.

Classification targets were based on liver biopsy evaluation. Three hepatic lesions (fibrosis (F0-F4), inflammation (I0-I5) and steatosis (S0-S3)) were dichotomized into three roughly balanced binary targets for classification (proportion of positive class is 54-56% in ≥F2, ≥I2, ≥S1 and 26% in ≥F3). Feature selection was performed first by ranking the proteins based on mutual information, then by a step-wise forward feature selection among the top 50 most important proteins to determine the optimal panel of features for each endpoint based on maximum ROC-AUC.

Data availability for the samples is unbalanced as not every endpoint neither every feature is available for each sample (patient). Therefore we split data to balance sets of patients by their pattern of availability for all considered variables. Each train and test split is based on this global missing pattern defined on the entire dataset allowing to compare subsets of patients with each other.

The metrics are compared on a test set which results from an 80% training and 20% test set split on the global pattern over the maximum of 360 samples in the high risk group, i.e. patient from which liver biopsies were taken. Each test set for each endpoint and marker combination is a subset of the globally defined test set on the 360 samples due to the imbalanced data.

The metrics for comparison are precision, recall, F1-score and balanced accuracy (Supplementary Tables 17-22). First, these are reported for fixed clinical cutoffs described in literature and used by clinicians. Fixed clinical cutoffs can be viewed as a threshold model, which is not fitted to specific data. Second, metrics on logistic regression models for markers based only on the marker itself as feature are reported.

The endpoints metrics are compared both using a cross-validation procedure (5-fold 10 times cross-validation) and a finally fitted model on which DeLong’s test was performed for the statistical comparison of AUCs. The cross-validation procedure is intended to give an overview on model performance variance from random data split. The final model is supposed to ease comparison with follow-up data on the patients in the high risk group and to have an explicit comparison based on one model. The final model was also used to perform rule-out validation in lower-incidence populations, which were not used for training. These include a healthy cohort and an at risk cohort. Their proteomics samples were preprocessed together with the samples for model training.

Harrell’s C-index was computed in the Stata software. In total 348 patients living in the Region of Southern Denmark were included in the computation. Follow-up data was retrieved from the electronic patient records at hospitals in Region of Southern Denmark, combined with the Danish National Registry of central personal identification numbers. We defined liver-related events as the occurrence of any of the following: alcoholic hepatitis, varices needing treatment (VNT), variceal bleeding, ascites, spontaneous bacterial peritonitis (SBP), hepatic encephalopathy (HE), hepatocellular carcinoma (HCC), hepatorenal syndrome (HRS), upper gastrointestinal bleeding, or jaundice due to liver failure.

## Data availability

All results from statistical and bioinformatics analysis were provided in the supplementary tables. All Python scripts can be reviewed and downloaded at the Github repository https://github.com/llniu/ALD-study.

## Acknowledgments

We are grateful to all patients and healthy volunteers who participated in the clinical cohort. We also thank all members of the Clinical Proteomics group and the Proteomics and Signal Transduction Group (Max Planck Institute). In particular, we thank Nicolai Jacob Wewer Albrechtsen for fruitful discussions, and Jeppe Madsen, Martin Rykær and Lylia Drici for technical assistance.

## Author contributions

L.N. designed experiments, performed and interpreted the MS-based proteomics analysis, carried out bioinformatics and machine learning analysis, and generated text and figures for the manuscript. M.T. designed the clinical cohort and provided insights in data analysis strategies. P.G., R.G., F.M. performed MS-based proteomics analysis. D.N.R. collected patient follow-up data, performed the prognostic performance analysis. H.E.W. performed machine learning analysis, generated corresponding Jupyter notebook. M.S. performed machine learning analysis. M.K., K.L., S.J. recruited participants in the clinical cohorts, collected samples and clinical data. A.S., S.R., T.H., A.K. revised the manuscript. M.M supervised and guided the project, interpreted MS-based proteomics data and wrote the manuscript.

