## Supplementary figures for "A paired liver biopsy and plasma proteomics study reveals circulating biomarkers for alcohol-related liver disease"

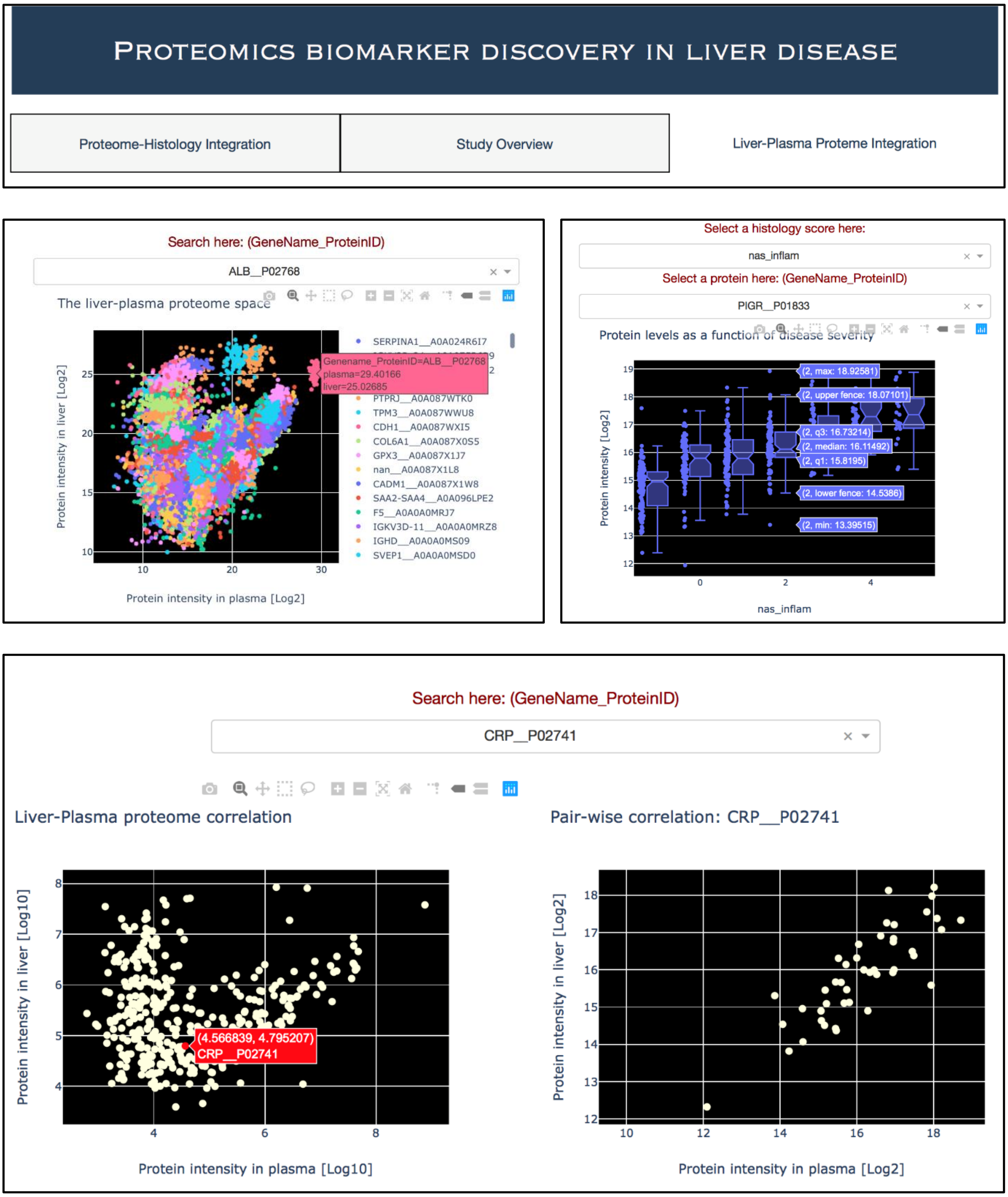


**Fig. S1 | An interactive, web-based data exploration app built with the open-source Dash framework.** The app enables dynamic data visualization and query for results, including protein abundance as a function of disease severity (liver histology scores), projection of plasma proteome to the liver space, and pairwise correlation between liver and plasma samples. Code and corresponding datasets will be provided in the Github repository https://github.com/llniu/ALD-study, and can be hosted on local machines.

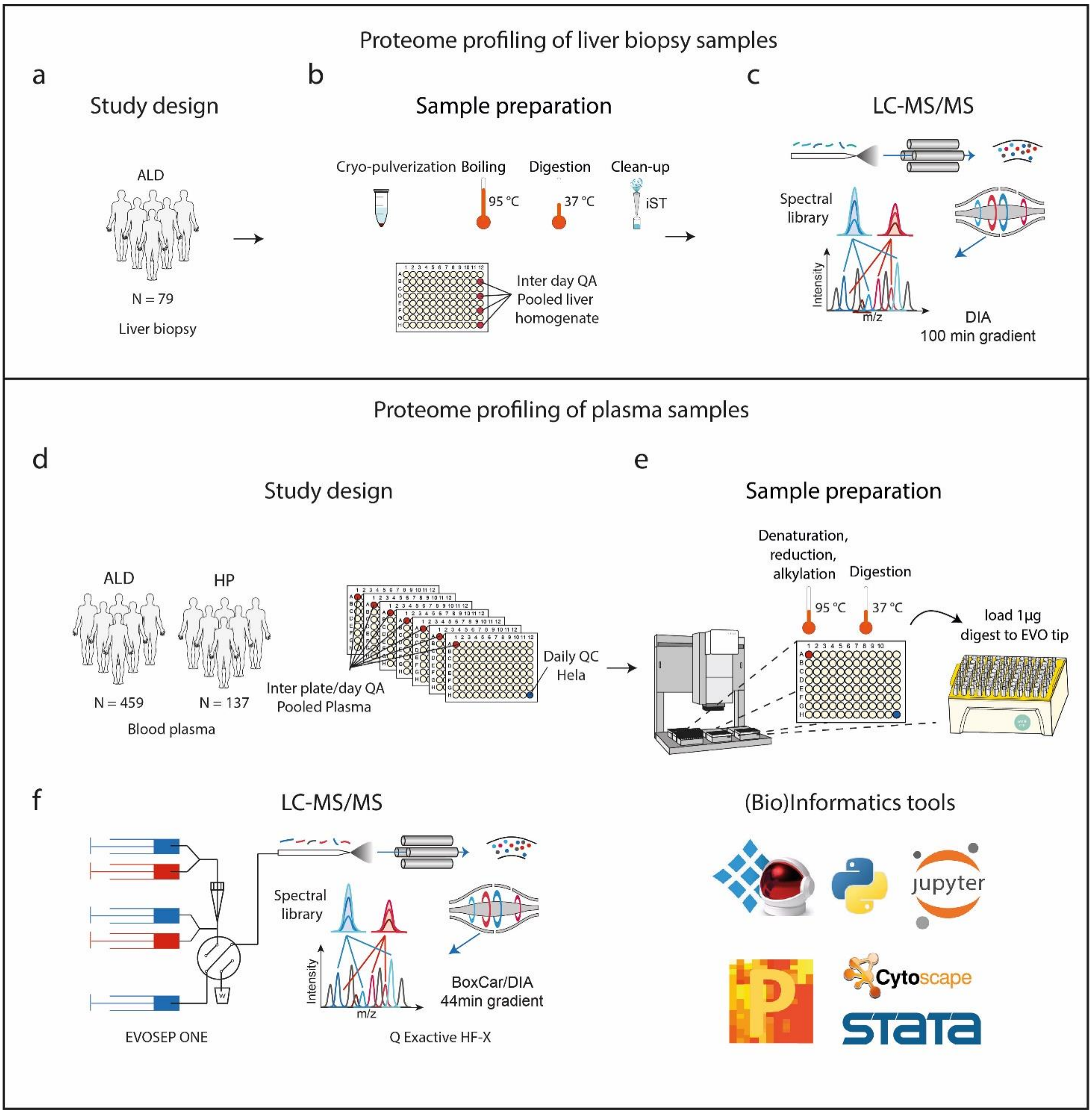


**Fig. S2 | Proteomics workflow. a-c.** Proteome profiling of liver biopsy samples. In total, 79 study samples were cryo-powderized, proteins were denatured, reduced, alkylated and digested in a 96-well plate using the PreOmics SDC lysis buffer peptides purified using the PreOmics iST protocol in a randomized manner. Purified peptides were analyzed by LC-MS/MS using a 100min-gradient (nanoEASY LC) with DIA acquisition mode in technical single shot. Four pooled homogenate samples were included as inter-day quality control samples to assess workflow variability from sample preparation to LC-MS/MS measurement. **d-f.** Proteome profiling of plasma samples. In total, 596 study samples and 7 pooled quality control samples were distributed in a randomized manner to 7 x 96-well plates. Proteins were denatured, reduced, alkylated and digested using the automated Plasma Proteome Profiling pipeline on the Bravo liquid handling system. Peptide digest were loaded to Evotips according to the EVOSEP tip-loading procedure. Peptide mixture were analyzed by LC-MS/MS using a 44-min gradient (EVOSEP One) with BoxCar/DIA acquisition mode in technical single shot. **g.** Resulting mass spectrometry raw files were processed by the Spectronaut software matching to a deep liver/plasma spectral library. Data analysis was performed in Python scripts in Jupyter Lab, the Persues software, CytoScape and the Stata software.

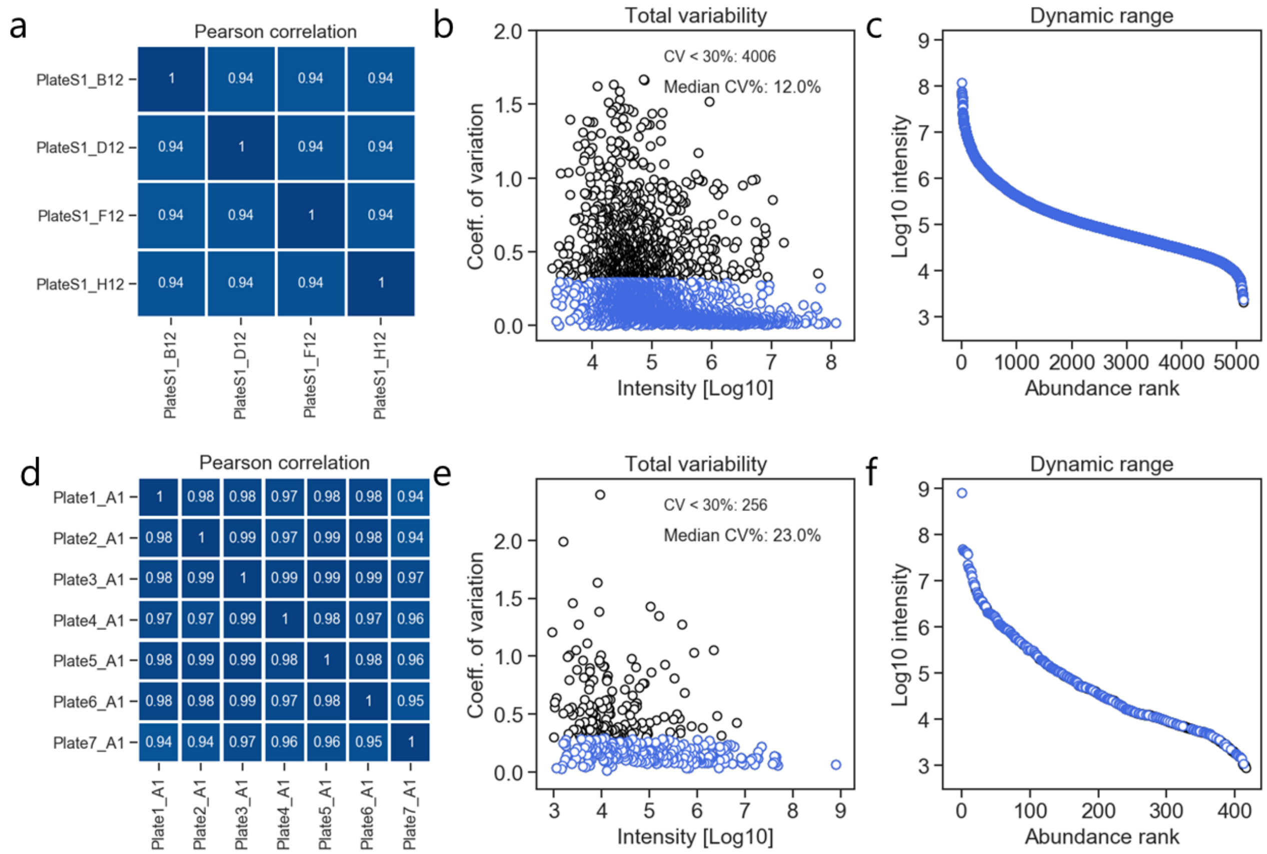


**Fig. S3 | Proteomics data quality. a,d.** Pair-wise Pearson correlation between proteomes of the workflow replicates (quality assessment samples, QA) in the liver (a) and plasma (d) proteomics experiment. **b, e**. Pearson coefficient of variation (CV) of each protein assessed by QA samples are plotted against their median intensity, with (b) showing the liver- and (e) showing the plasma proteomics experiment. Proteins shaded in blue have a CV% < 30%. **c, f.** Protein intensity as a function of abundance rank in the liver-and plasma proteomes (c and f, respectively).

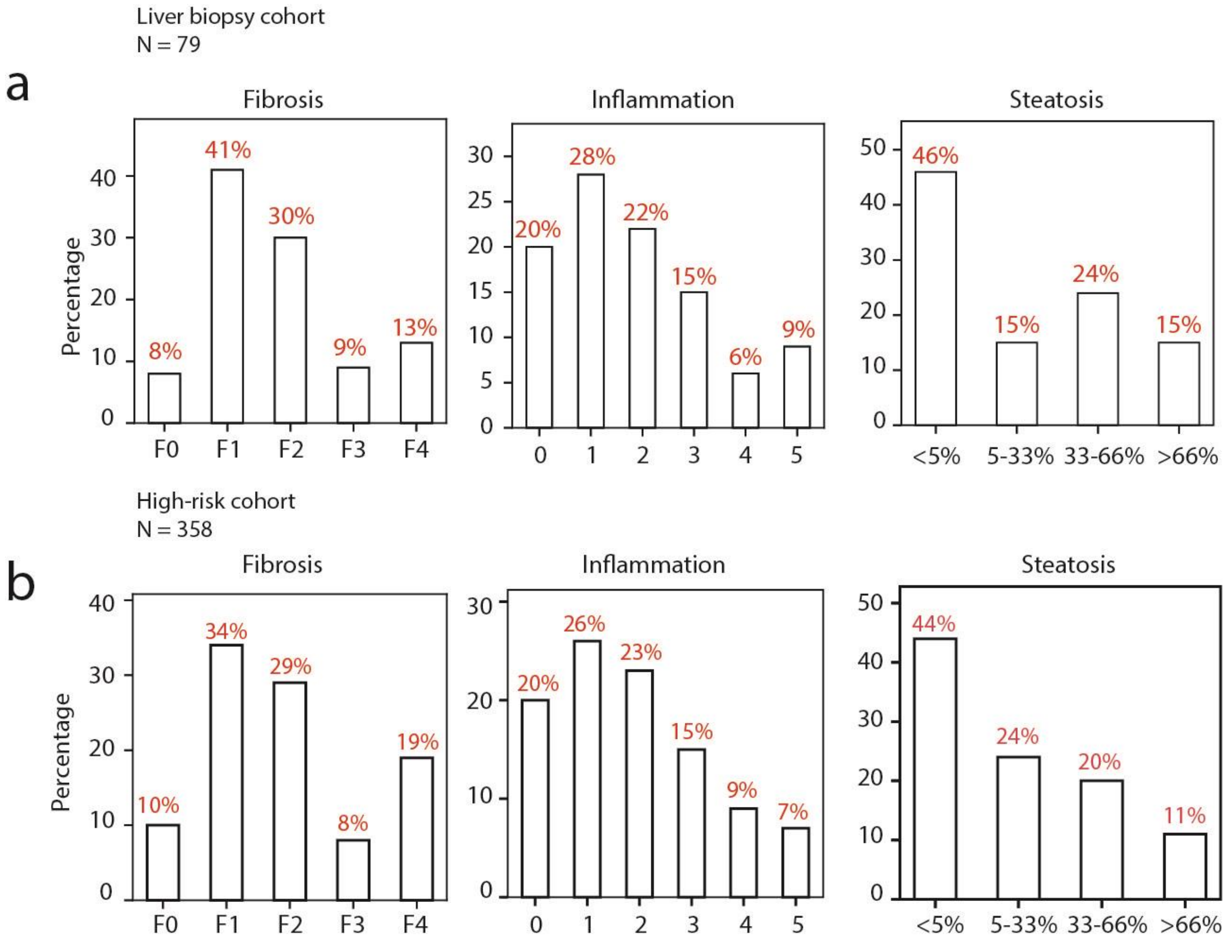


**Fig. S4 | Histologic score distribution in the high-risk cohort. a.** Stage distribution of fibrosis, inflammation and steatosis in patients whose liver biopsy proteomes were analyzed. Total number of samples are noted. **b.** Stage distribution of fibrosis, inflammation and steatosis in patients whose plasma proteome were analyzed.

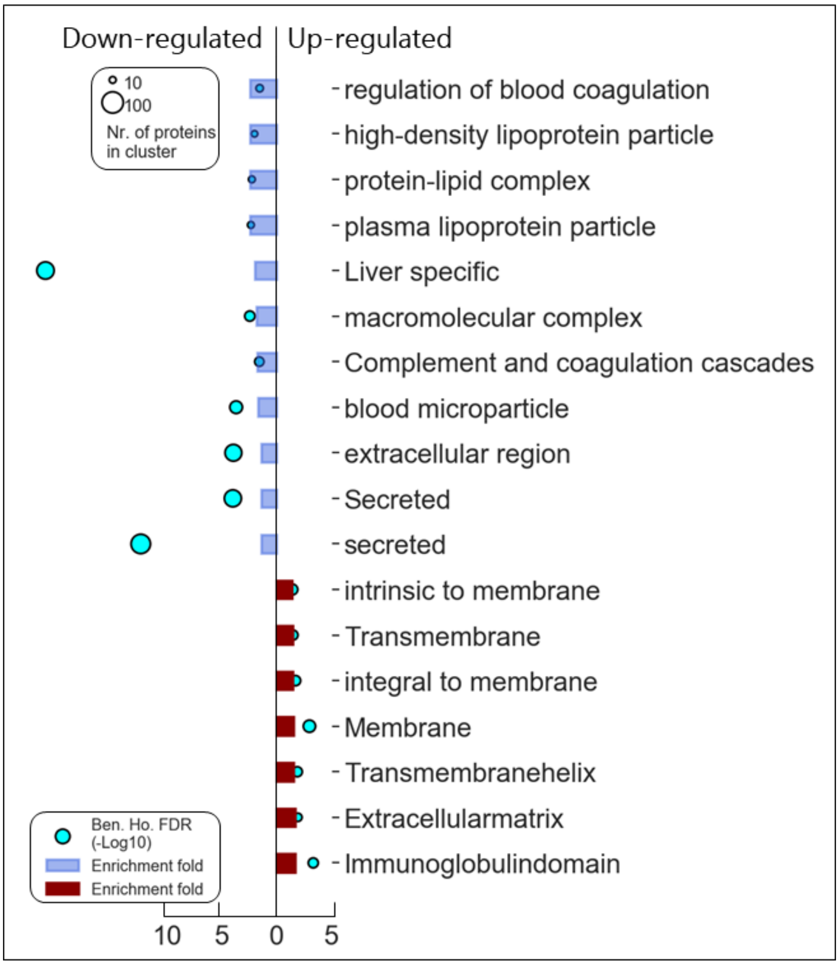


**Fig. S5 | Functional enrichment in plasma proteome.** Significantly enriched terms in the down- and upregulated plasma proteins as liver fibrosis progresses are shown, with enrichment fold and Benjamini-Hochberg corrected FDR annotated. Circle sizes indicate number of proteins involved in each term.

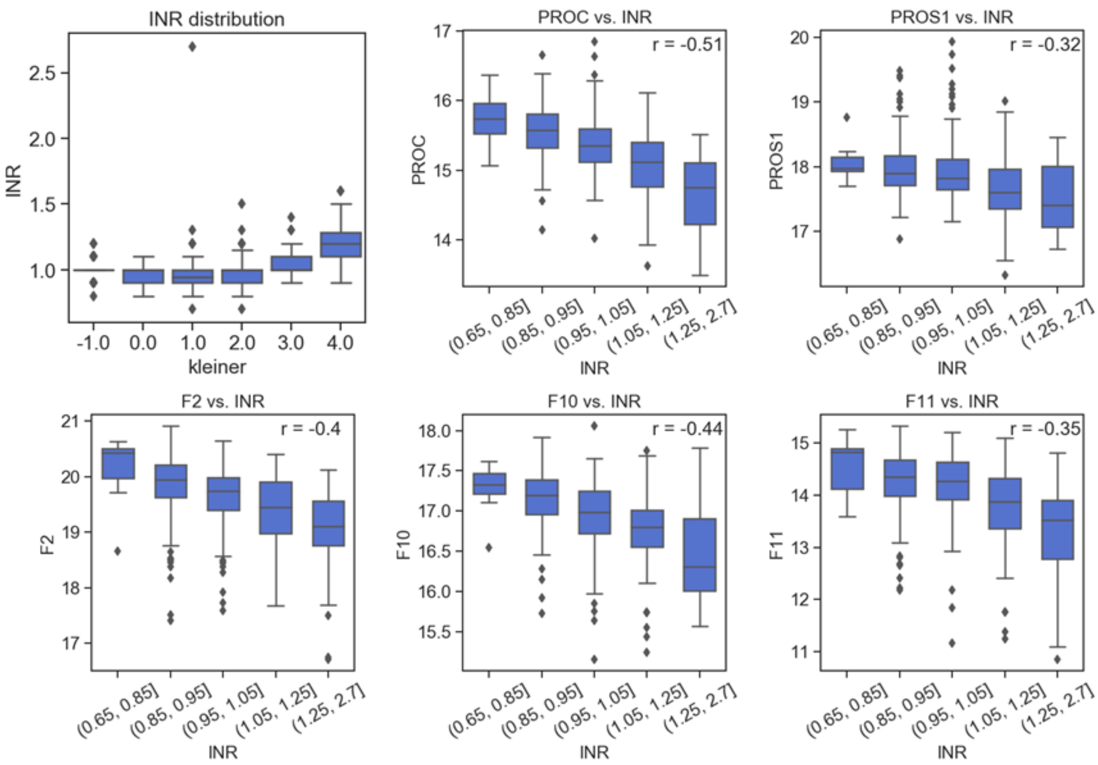


**Fig. S6 | INR distribution across fibrosis stages and Pearson correlation between clotting factors (PROC, PROS1, F2, F10, F11) and INR.**

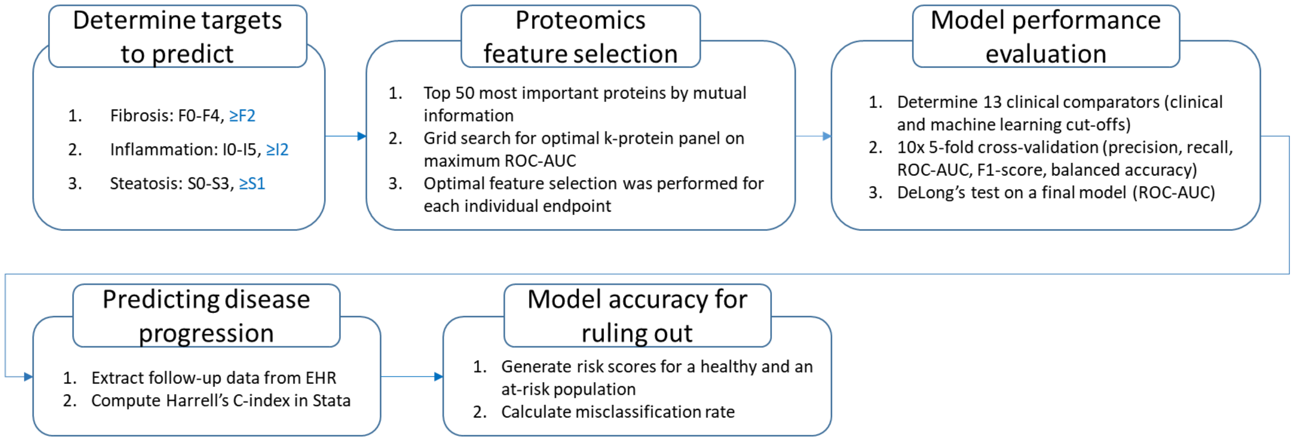


**Fig. S7 | Overview of machine learning pipeline.** Graphic workflow detailing the target binary classification problems, feature selection strategies, model performance evaluation strategies, prognostic power assessment and validation of model accuracy for ruling out liver injury in low-incidence populations.
